# FAM122A inhibition of PP2A-B55 through a bipartite binding mechanism

**DOI:** 10.64898/2026.03.24.713894

**Authors:** Iker Benavides-Puy, Suzanne Vigneron, Arminja N. Kettenbach, Thierry Lorca, Jakob Nilsson

## Abstract

FAM122A regulates cell cycle progression through inhibition of the PP2A-B55 phosphoprotein phosphatase. Recent structural work has uncovered helical elements in the N-terminus of FAM122A as binding determinants for PP2A-B55 but whether FAM122A inhibition towards PP2A-B55 is regulated is presently unclear. To address this we performed a systematic analysis of the PP2A-B55 interaction with FAM122A in cells uncovering a novel region in the C-terminus of FAM122A, spanning residues 150-170, required for binding. This C-terminal region and the N-terminal helices are both required for efficient binding to PP2A-B55 suggesting a bipartite binding mechanism. We perform amino acid resolution scans of FAM122A 150-170 uncovering several residues in this region contributing to binding including the conserved Ser158, a reported phosphorylation site. We show that Ser158 is important for PP2A-B55 inhibition in human cells as well as efficient stimulation of mitotic entry in *Xenopus laevis* egg extracts. In human cells and in *Xenopus laevis* Ser158 phosphorylation is regulated with increased occupancy correlating with cell cycle stages requiring PP2A-B55 inhibition. Collectively our work uncovers novel aspects of FAM122A interaction with PP2A-B55 and provides a possible mechanism for how the inhibitory activity of FAM122A can be regulated during the cell cycle.

## Introduction

The Ser/Thr phosphoprotein phosphatase PP2A-B55 is a major cell cycle regulator and composed of a catalytic subunit (PPP2CA), a scaffold subunit (PPP2R1A/B) and the B55 regulatory subunits (B55α-δ) providing substrate specificity (*1–3*). PP2A-B55 regulates the cell cycle by antagonizing the activity of cyclin dependent kinases (CDKs). For efficient mitotic entry it is critical that the activity of PP2A-B55 is inhibited allowing unrestrained CDK phosphorylation of thousands of proteins. To achieve temporal PP2A-B55 inhibition CDK1/Cyclin B1 activates the Greatwall (Gwl)/MASTL kinase that phosphorylates two small unstructured proteins, ARPP19 and ENSA (*4–10*). ARPP19 and ENSA are very similar in sequence and their inhibition of PP2A-B55 depends on the phosphorylation by Gwl/MASTL of a conserved Ser residue (Ser62/Ser67 respectively). Recent cryo-EM structures of thiophosphorylated ARPP19 bound to PP2A-B55 shows that ARPP19 positions the phosphorylated Ser62 in the active site of PPP2CA while N-terminal helical elements bind the substrate binding pocket on B55 (*11, 12*). In addition to ARPP19/ENSA another small protein, FAM122A, has recently been reported to act as an inhibitor of PP2A-B55 to control the cell cycle and the response to Chk1 inhibition (*13–15*). The cryo-EM structure of a truncated FAM122A protein bound to PP2A-B55 shows that two N-terminal helices in FAM122A binds B55 and blocks the catalytic site respectively (*11*). However, work in *Xenopus laevis* egg extract suggests that like ARPP19/ENSA the inhibitory activity of FAM122A might be stimulated by phosphorylation although the exact phosphorylation site(s) involved in this are unclear (*14*). To increase our understanding of FAM122A as an inhibitor of PP2A-B55 we here conduct a systematic analysis of the full-length protein in cells to uncover a conserved region in the C-terminus of the protein required for PP2A-B55 binding and inhibition. This region contains a highly conserved Ser residue that is critical for FAM122A function, and we show that it is phosphorylated upon mitotic entry in *Xenopus laevis* suggesting that FAM122A inhibitory activity towards PP2A-B55 might be cell cycle regulated.

## Results

### FAM122A is an inhibitor of PP2A-B55 acitivity in cells

Recent work has uncovered that FAM122A is an inhibitor of PP2A-B55 and that this inhibition can contribute to cell cycle regulation. Whether FAM122A inhibitory activity towards PP2A-B55 is regulated is not fully understood. To investigate this we first wanted to confirmed that FAM122A is an inhibitor of PP2A-B55 in human cells. To do this we transfected HeLa cells with YFP or YFP-FAM122A and 24 hours later we immunopurified PP2A-B55 from asynchronous cells and measured phosphatase activity towards a model peptide substrate (**Fig. 1A**)(*16*). The expression of YFP-FAM122A reduced PP2A-B55 activity by ∼40% in line with its reported inhibitory activity. We next generated a FAM122A knockout U2OS/FRT/TRex cell line using CRISPR-Cas9 technology (**Fig. 1B**) and analysed the sensitivity of this cell line to low doses of okadaic acid (OA). Okadaic acid is a broad spectrum inhibitor of phosphoprotein phosphatases but at low nanomolar concentrations it largely affects PP2A, PP4 and PP6 (*17*). The knockout of FAM122A resulted in decreased sensitivity to OA suggesting higher levels of cellular PPP activity in the absence of FAM122A (**Fig. 1C**). This effect was not due to changes in cell cycle distribution (**Fig. S1A**). To confirm that this effect was due to the absence of FAM122A we reintroduced YFP-FAM122A using the FRT system under a doxycyclin inducible promoter. The induction of YFP-FAM122A expression increased sensitivity to OA confirming that the observed effects in the U2OS FAM122A knockout cell line is indeed due to the absence of FAM122A. Based on the selective binding of FAM122A to PP2A-B55 we favour that the decreased sensitivity to OA upon FAM122A removal is caused by increased PP2A-B55 activity. Collectively these experiments support that in cells FAM122A is an inhibitor of PP2A-B55.

**Fig. 1:**
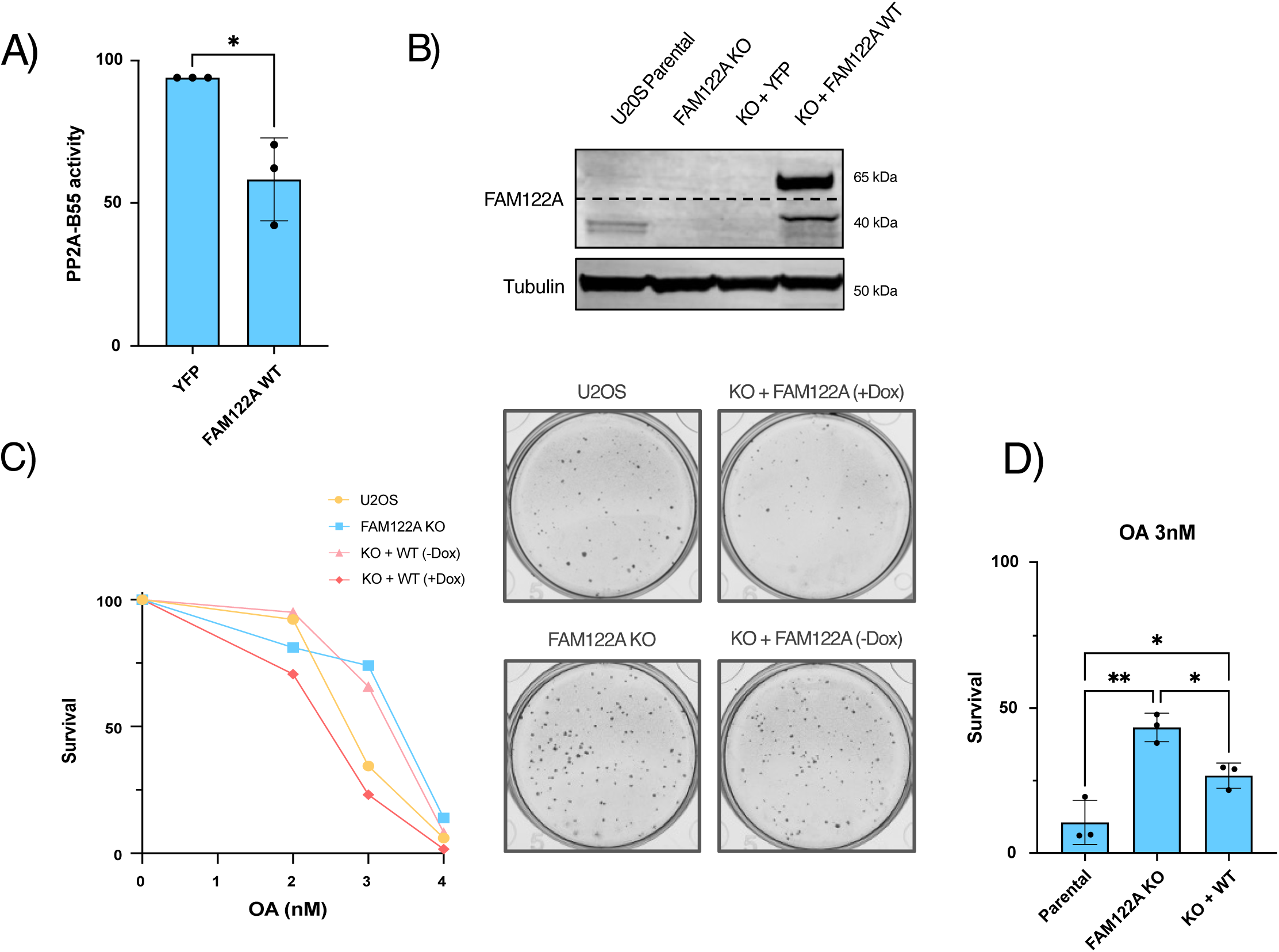
FAM122A is an inhibitor of PP2A-B55. **A)** The indicated YFP constructs were expressed in HeLa cells, followed by PP2A-B55 immunoprecipitation, and subsequent phosphatase activity assay using a model peptide substrate. Quantification shows the average and standard deviations of 3 biological replicates. **B)** Immunoblotting of the indicated stable cell lines showing the levels of FAM122A and tubulin. **C)** Colony formation assay performed in the indicated U2OS/FRT/Trex cell lines showing sensitivity to different concentrations of OA. The stable FAM122A WT cell lines were treated with 10 ng/mL of doxycycline or untreated. Representative images for 3 nM OA are shown. **D)** Bar plot showing sensitivity of the cell lines to 3nM of OA. Quantification shows the average and standard deviations of 3 biological replicates.

### Efficient binding of FAM122A to PP2A-B55 in cells requires N-terminal and C-terminal elements

The cryo-EM structure of FAM122A 1-124 bound to PP2A-B55 revealed two helixes in the region 81-111 making direct contact to PP2A-B55 (*11*). One helix binds the substrate binding pocket of B55 while the second helix binds the catalytic subunit to inhibit phosphatase activity. These observations where also supported by parallel studies using AlphaFold modelling and mutational analysis (**Fig. 2A**) (*12, 14, 15*). When we analyzed a truncation panel of FAM122A for the ability to bind PP2A-B55 in HeLa cells we were surprised to find that efficient binding required sequence elements in the 150-200 region of FAM122A in addition to the N-terminal region (**Fig. 2B**). To further confirm this we conducted a systematic analysis of this region. A 10 amino acid truncation analysis of this region revealed that amino acids from 150-170 was required for binding PP2A-B55 (**Fig. S1B**). This conclusion was supported by a 10 amino acid Ala scan through this region (**Fig. 2C**).

**Fig. 2:**
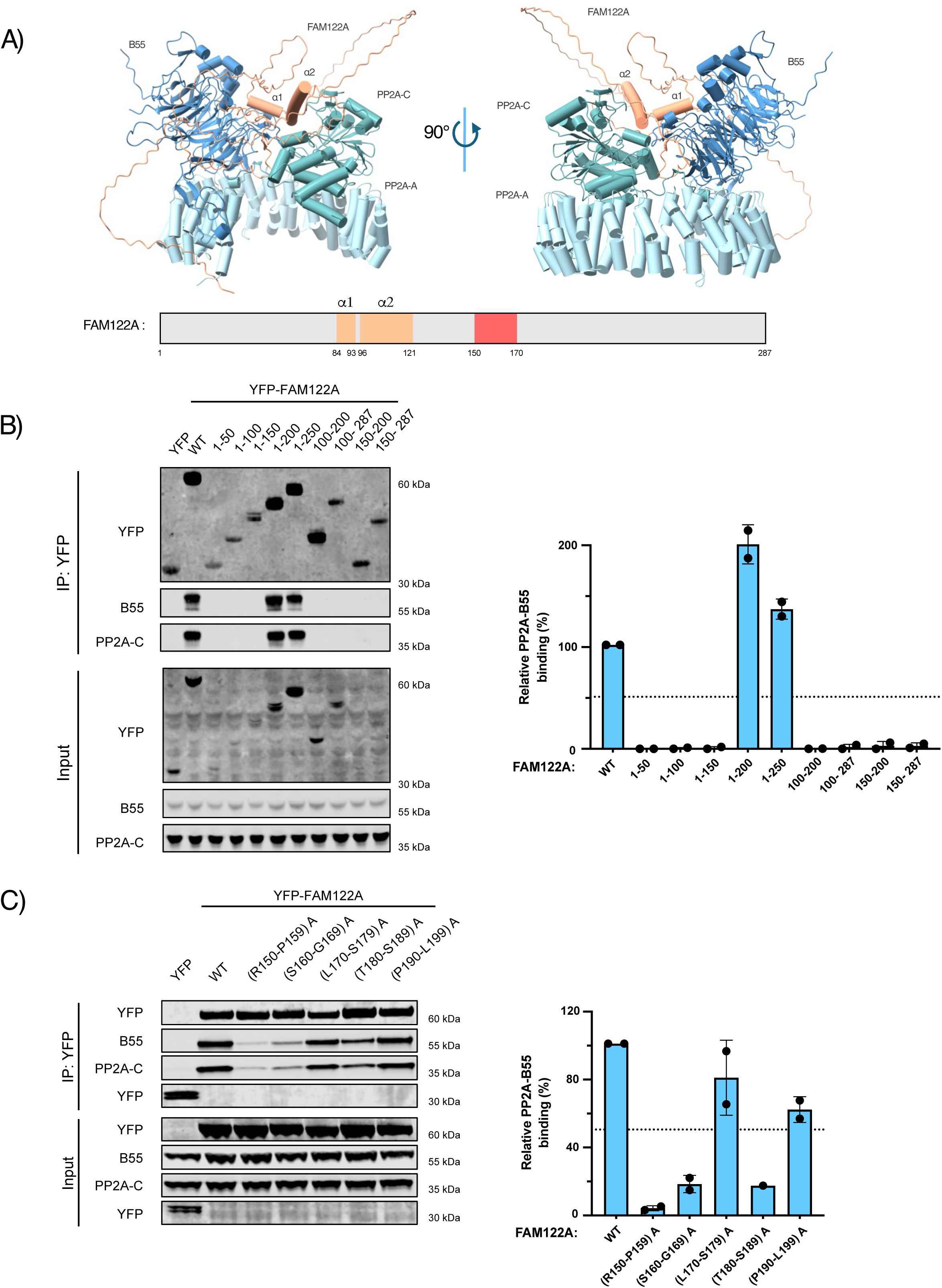
N-terminal and C-terminal binding determinants in FAM122A. **A)** AlphaFold3 model of full-length FAM122A in complex with PP2A-B55α. Below, a schematic representation of FAM122A, highlighting the PP2A-B55 binding helices, and in red the region encompassing amino acids 150-170. **B-C)** The indicated YFP-tagged FAM122A fragments or mutants were expressed in HeLa cells, followed by YFP trap pull down and subsequent immunoblotting of indicated proteins: PP2A-C; PP2A catalytic subunit; PP2A-B55, B55 regulatory subunit, and YFP. Quantification of two biological replicates (showing the average and standard deviations) assessing relative binding of FAM122A constructs indicated to PP2A-B55 normalized to WT protein.

To identify the exact residues in FAM122A 150-170 contributing to binding we conducted a single alanine scan through this region. Several residues throughout this region contributed to FAM122A-PP2A-B55 interaction but all residues in the region spanning residues 154-159 where required for efficient binding (**Fig. 3A-B**). This region of FAM122A is highly conserved despite being unstructured (**Fig. 3C**).

**Fig. 3:**
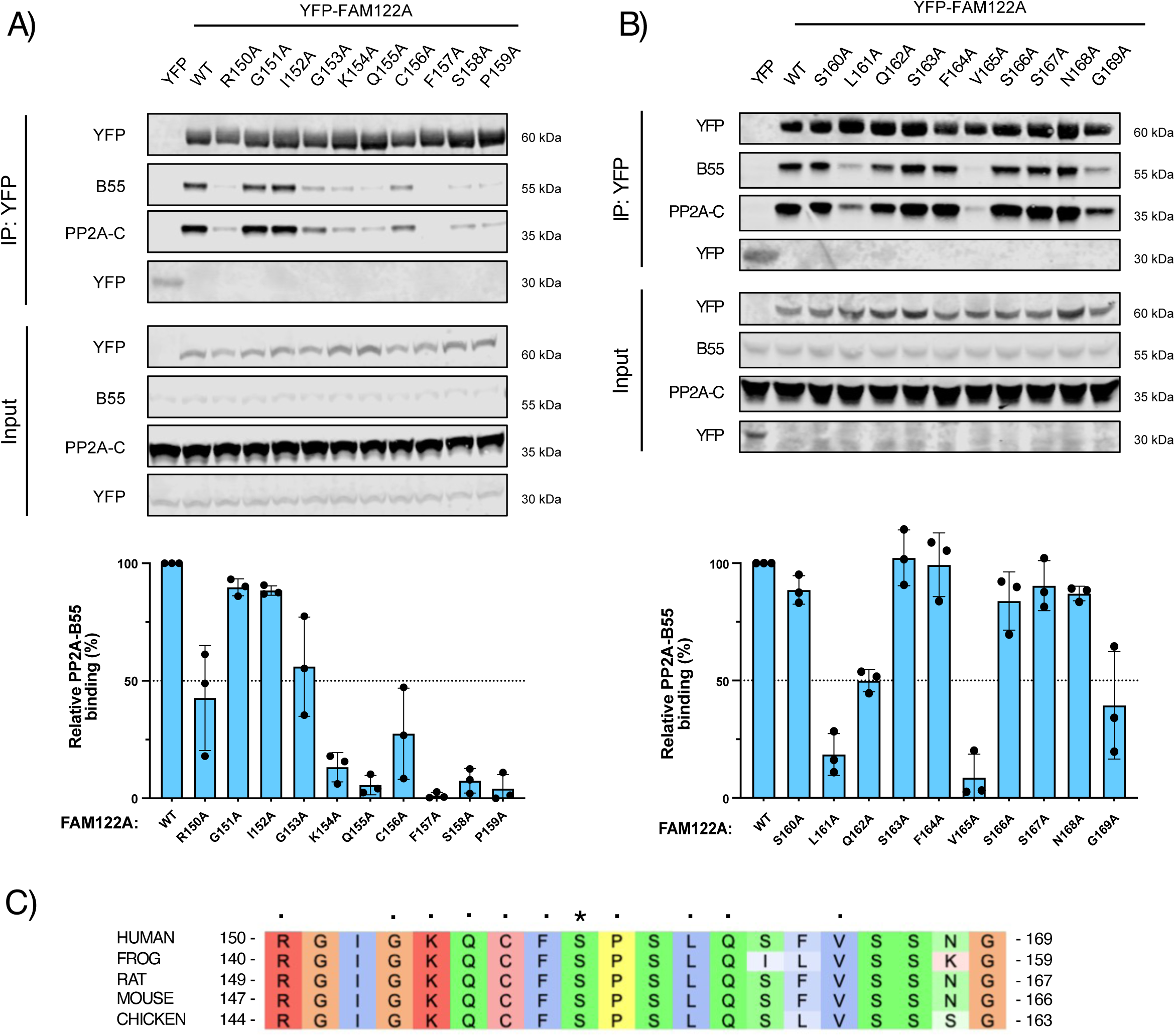
Single alanine scan through 150-170 identifies critical residues for PP2A-B55 binding. **A-B)** The indicated YFP-FAM122A single alanine mutants were expressed in HeLa cells, followed by YFP trap pull down and subsequent immunoblotting of indicated proteins: PP2A-C; PP2A catalytic subunit; PP2A-B55, PP2A regulatory subunit, and YFP. Quantification of three biological replicates (showing the average and standard deviations) assessing relative binding of FAM122A constructs indicated to PP2A-B55 normalized to WT protein. **C**) Alignment of 150-170 region from FAM122A proteins from different species, with residues critical for binding indicated with a •, and reported phosphorylation sites with an *.

We conclude that in cells efficient FAM122A-PP2A-B55 complex formation requires the N-terminal helices as well as several residues in the region spanning 150-170. As neither the N-terminus nor C-terminus alone binds efficiently in cells this suggests a bipartite binding mechanism.

### FAM122A Ser158 is required for PP2A-B55 inhibition and efficient mitotic entry in *Xenopus laevis*

The FAM122A 154-159 region required for PP2A-B55 binding encompasses Ser158 a reported phosphorylation site (phosphosite.org). This phosphorylation site matches a proline-directed kinase consensus sequence suggesting that it could be phosphorylated by CDKs and that this might regulate FAM122A binding or inhibition of PP2A-B55 during the cell cycle. The FAM122A S158A mutation prevented binding to all isoforms of PP2A-B55 (**Fig. 4A and Table 1**). We tested the ability of FAM122A S158A to inhibit PP2A-B55 as described above and included a FAM122A mutant with the B55 binding helix mutated (RL-IK mutant). Both of these mutants did not inhibit PP2A-B55 phosphatase activity (**Fig. 4B and S2A**). AlphaFold 3 modelling (*18*) of full length FAM122A with or without Ser158 phosphorylation modelled a movement of this region upon phosphorylation with the phosphate group occupying the active site of PP2AC (**Fig. 4C**). This binding of the phosphate group to the active site did not appear to require a major movement of the helix observed in the cryo-EM structure to bind the catalytic subunit. These models promted us to further explore FAM122A Ser158 phosphorylation

**Fig. 4:**
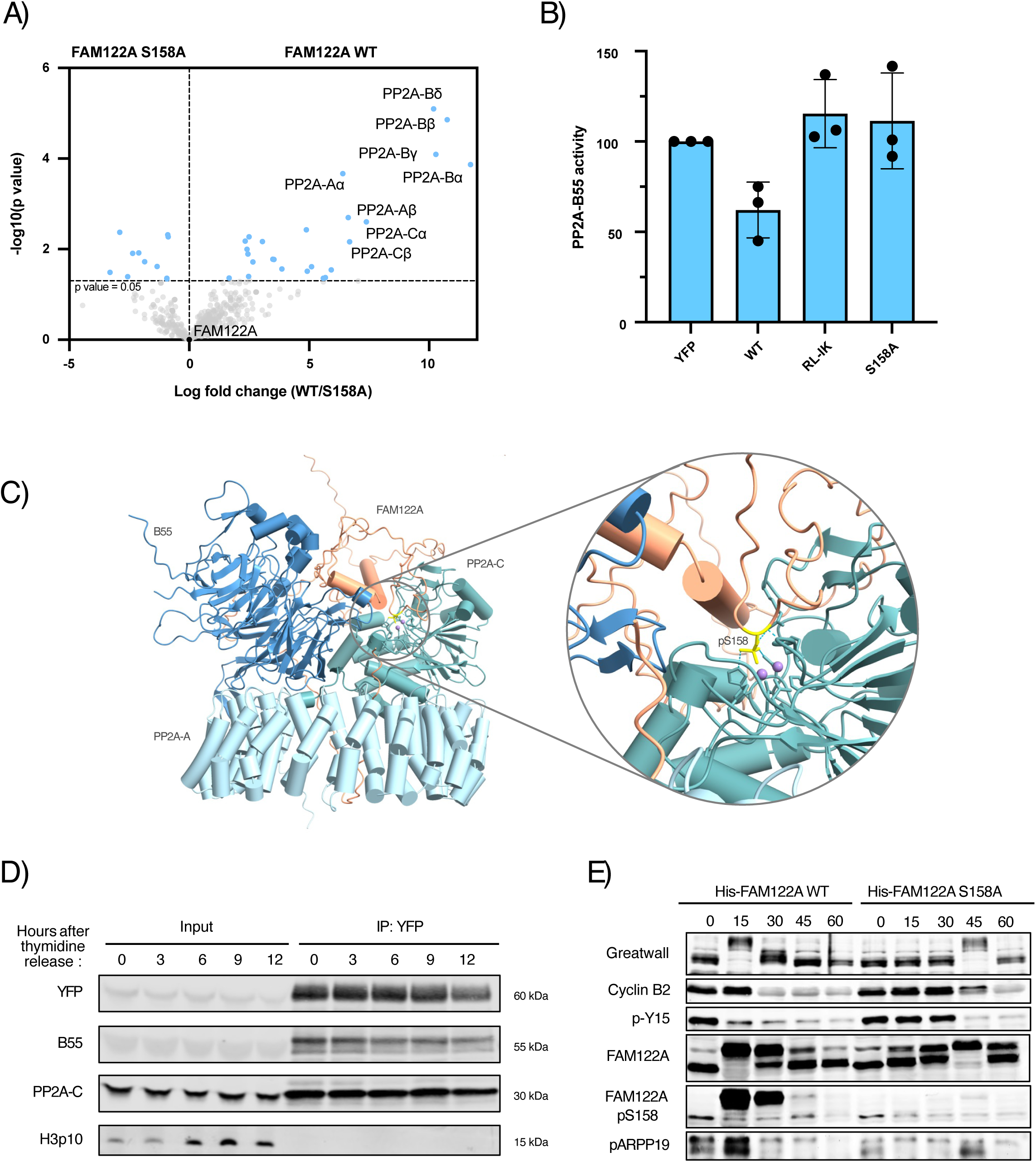
FAM122A Ser158 is required for PP2A-B55 inhibition and efficient mitotic entry. **A)** YFP-tagged FAM122A WT or FAM122A S158A constructs were expressed in asynchronous HeLa cells followed by YFP trap pull down and mass spectrometry analysis of three technical replicates (PP2A-Bα; B55 alpha subunit, PP2A-Bβ; B55 beta subunit, PP2A-B55δ; B55 delta subunit, PP2A-Bγ; B55 gamma subunit, PP2A-Cα; catalytic subunit alfa, PP2A-Cβ; catalytic subunit beta, PP2A-Aα; scaffold subunit alfa, PP2A-Aβ; scaffold subunit beta). **B)** The indicated YFP-FAM122A constructs were expressed in HeLa cells, followed by PP2A-B55 immunopurification, and subsequent phosphatase activity assay using a model peptide. RL-IK mutant 84-**RL**HQ**IK**QEE-92. Quantification showing the average and standard deviation of 3 biological replicates assessing PP2A/B55 activity is shown. **C)** AlphaFold3 model of full-length FAM122A S158p in complex with PP2A-B55 highlighting the positioning of the phosphate group in the active site. **D)** YFP-FAM122A WT was expressed in HeLa cells and synchronized by double Thymidine release. Cells were released at T_0_, and collected 3, 6, 9, and 12 hours after release, followed by GFP trap pull down and subsequent immunoblotting of indicated proteins: PP2A-C; PP2A catalytic subunit; PP2A-B55, B55 regulatory subunit, H3p10, phosphorylation on S10 in Histone 3, and YFP. Representative blot of three biological replicates. **E)** 20 µl of interphase extracts was supplemented with 2 µl (0.66 µg) of either the wild-type Human FAM122A or 3 µl (0.66µg) of S158A FAM122A mutant and 1 µl of the mix was recovered at different time points to follow the indicated proteins by western blot. Representative of three biological replicates.

**Table 1:** Analysis of YFP-FAM122A purifications by mass spectrometry.

To analyse regulation of Ser158 phosphorylation we purified YFP-FAM122A from asynchrounous and mitotic cells and performed a quantitative mass spectrometry analysis. A peptide encompasing Ser158 and Ser172 revealed that the double phosphorylated peptide was slightly more abundant in the mitotic sample while the single phosphorylated peptides showed a modest decrease in the mitotic samples (**Fig. S2B**). This shows that Ser158 has a higher occupancy of phosphorylation during mitosis. Since Ser158Ala, Ser158Glu and Ser158Thr mutations all prevent binding to PP2A-B55 in cells this argues that phosphorylation of Ser158 would impact binding but we cannot establish if this would be a positive or negative effect as the mutation to Ala or Glu both prevent binding (**Fig. S2C**). Analysis of FAM122A-PP2A-B55 interaction during progression from S-phase to M-phase in HeLa cells revealed no major changes in interaction arguing that the Ser158 phosphorylation might not affect binding during an unpertubed cell cycle in HeLa cells (**Fig. 4D**).

To further explore a potential function of Ser158 and its phosphorylation we turned to the *Xenopus laevis* egg extract system that allows high temporal resolution of mitotic entry. Given the high temporal resolution of the system we deemed it more suited to explore regulatory mechanisms of FAM122A. Previous work has shown that addition of recombinant full length human FAM122A to interphase extracts stimulates mitotic entry and that a mutant version of FAM122A with all Ser/Thr sites mutated to Ala is less efficient to promote mitotic entry (*14*). This phosphonull mutant seems proficient in binding PP2A-B55 arguing that in this system the Ser158Ala mutation does not affect binding (*14*). To probe functionality of Ser158 we analysed mitotic entry by western blot following addition of full length recombinant human FAM122A WT and Ser158Ala to interphase extracts (**Fig. 4E**). We observed a delay of 30 minutes in Gwl hyperphosphorylation, disapperance of Cyclin B2 and Cdk1 Y15 phosphorylation in FAM122A Ser158Ala complemented extracts consistent with a delay in mitotic entry. Analysis of FAM122A migration revealed retardation of migration following 15 minutes of incubation and this was delayed in the Ser158Ala mutant. Indeed a phosphospecific FAM122A Ser158 antibody revealed strong phosphorylation of FAM122A at the earliest time points followed by dephosphorylation as cells exited mitosis.

## Discussion

PP2A-B55 is an important regulator of the cell cycle due to its ability to dephosphorylate CDK phosphorylation sites (*12, 16, 19, 20*). At least three PP2A-B55 specific inhibitors (ARPP19, ENSA and FAM122A) control phosphatase activity and cell cycle progression. In the case of ARPP19 and ENSA this inhibitory activity is regulated by phosphorylation while the current structural knowledge of FAM122A-PP2A-B55 does not provide clues to potential regulation. From a biological perspective it seems advantageous to be able to regulate FAM122A inhibition of PP2A-B55 to control signaling pathways.

The data presented here provides an important step towards understanding FAM122A regulation by the identification of Ser158 as a critical binding determinant for efficient PP2A-B55 binding and inhibition. This conclusion is supported by the inability of FAM122A Ser158Ala to efficiently promote mitotic entry in *Xenopus laevis* egg extracts. We show that Ser158 phosphorylation is regulated during the cell cycle and this phosphorylation could be mediated by CDKs given the higher occupancy in mitosis. It is also highly likely that other proline directed kinases can phosphorylate FAM122A Ser158 in response to specific cellular perturbations to regulate PP2A-B55 activity. Identifying these kinases and pathways will be important future directions. The exact function of Ser158 phosphorylation is at present unclear to us as both Ser158Ala and Ser158Glu are unable to bind PP2A-B55 either reflecting that phosphorylation is inhibitory or that Glu is not a true phosphomimetic mutation. We favor the later possibility since in most cases where a phosphorylation has a unique function in binding the phosphomimetic mutations normally do not recapitulate this (*21*). Indeed, phosphomimetic variants of ARPP19 and ENSA are not effective inhibitors of PP2A-B55. Given the rapid phosphorylation of FAM122A Ser158 prior to mitotic entry in *Xenopus laevis*, a time where PP2A-B55 needs to be inhibited, this could be suggestive of Ser158 phosphorylation positively regulating FAM122A inhibition of PP2A-B55. Our AlphaFold3 modelling suggests that S158 phosphorylation could potentiate PP2A-B55 inhibition by positioning the phosphate group in the active site of PP2AC similar to how ARPP19 and ENSA inhibits this phosphatase. We speculate that the reason why FAM122A Ser158Thr is unable to bind PP2A-B55 is due to more efficient dephosphorylation of Thr residues by PP2A-B55 and that FAM122A is both and inhibitor and substrate of the phosphatase similar to the described mechanism for ARPP19/ENSA (*22, 23*).

## Acknowledgments

Work at the Center for Epigenetic Cell Memory, Danish Cancer Institute is supported by The Danish National Research Foundation (DNRF195). Work at the Novo Nordisk Foundation Center for Protein Research is supported by NNF14CC0001. Work in the Nilsson lab is supported by grants from the Novo Nordisk Foundation (NNF0082227 and NNF0065098), DFF-FNU (4283-00188B) and the Danish Cancer Society (R269-A15586-B71). This work was supported by grant R35GM119455 from the National Institute of General Medicine to ANK and the NCI Cancer Center Support Grant P30CA023108. We would like to thank Marc Plays and Anthony Ollier for providing the Xenopus females, the ZEFIX animal facility and for immunising the rabbits and recovering the antibodies. The work in the laboratory is supported by grants from the Agence Nationale pour la Recherche (ANR, France – ANR-22-CE13-0022), and the Fondation ARC pour la Recherche sur le Cancer (ARCPJA2025080010450).

## Materials and Methods

### Cell culture

HeLa cells (ATCC), U2OS Flp-In T-REx cells, and U2OS-derived cell lines were cultured in Dulbecco’s Modified Eagle Medium (DMEM) with GlutaMAX (Life Technologies) supplemented with 10% fetal bovine serum (Gibco) and Penicillin-Streptomycin (P/S) 10.000 units/mL (Invitrogen) at 37 °C with 5% CO2. The expression of the constructs in the U2OS Flp-In T-REx derived cell lines was induced upon treatment with 10 ng/mL of doxycycline (Clontech) for 24 hours unless otherwise stated.

### FAM122A gRNA cloning

The sequence for the gRNA against FAM122A was selected from the Brunello library. Forward and Reverse primers for the KO were designed adding to the gRNA the restriction sites for BbsI restriction enzyme. pX459 vector was digested with BbsI restriction enzyme and the product was gel purified.

gRNA for FAM122A KO:

5’ GCA-GAT-AAG-CCA-CTC-CTG-GG 3’

5’ CGT-CTA-TTC-GGT-GAG-GAC-CC 3’

### Generation of FAM122A KO U2OS Flp-In T-REx cell line

U2OS Flp-In T-REx cells were grown in DMEM supplemented with FBS and P/S. Cells were transfected with the pX459-FAM122A gRNA vector using the jetOPTIMUS (polyplus) reagent. After 24 hours of incubation, cells were split into 10 cm dishes at different dilutions in medium supplemented with 10% FBS, P/S, and 1 µg/mL of puromycin. After 48 hours, the medium was changed to a medium supplemented with Blasticidin and Zeocin. After 5 days, a confluent dish was collected, and screened by WB for FAM122A levels. 24 single clones were analysed by WB to confirm FAM122A knockout.

### Making FAM122A constructs stable cell lines in Flp-In T-Rex cells

U2OS Flp-In T-Rex FAM122A KO cells were seeded and transfected with pOG44 and pcDNA5 for each FAM122A construct, using jetOPTIMUS (polyplus). After 24 hours of incubation, and every 4-5 days until clear colonies were formed, the medium was replaced with DMEM + FBS + P/S supplemented with Blasticidin and Hygromycin. After clear colonies were formed, cells were collected and FAM122A expression was validated by WB.

### Transfection

For transient protein expression, cells were transfected with 2 µg of DNA of the indicated construct using jetOPTIMUS (polyplus) transfection reagent and incubated for 24 hours before collecting.

### Colony formation assay (CFA)

U2OS Flp-In T-REx parental cells or FAM122A KO cells stably expressing YFP-FAM122A WT cells were trypsinized and reseeded into six-well plates. 200 cells for each condition were seeded into a well. After seeding into the six-well dish, YFP-FAM122A WT cells were treated with the indicated concentration of Doxycycline. After 24 hours, cells were treated with the corresponding concentration of OA. After 7 days, cells were washed once in PBS, fixed, and stained in 0.5% methylviolet + 25% methanol. The solution was removed after incubation, and two washes with water were performed. Once they were dry, the plates were scanned on a GelCount (Oxford Optronix), and the number of colonies was quantified using the GelCount software. Each well’s plating efficiency (%) was calculated as the number of colonies divided by the number of cells seeded times 100. The surviving fraction for each dose of drug was calculated by normalizing the plating efficiency of the treated condition to the untreated one. Graphs were made in PRISM, and one-way ANOVA analyses with multiple comparison tests were performed comparing the mean of each condition to the parental cell line.

### Antibodies

Commercially available antibodies against the following proteins were used for western blotting in the indicated dilutions:

|  | Brand/reference | Catalog Number | Made in | Dilution |
| --- | --- | --- | --- | --- |
| PP2A-B55α | Cell Signaling Technology | 5689S | Mouse | 1:2000 |
| PP2A-C $\alpha/\beta$ | Merck | MABE1783 | Mouse | 1:2000 |
| GFP | Roche | 11814460001 | Rabbit | 1:5000 |
| FAM122A | Invitrogen | MA5-24510 | Mouse | 1:500 |
| FAM122A S158p | This study | - | Rabbit | 1:500 |
| Tubulin | Abcam | Ab6160 | Rat | 1:10000 |
| Xenopus Greatwall | Vigneron et al. 2009(8) | - | Rabbit | 5 ug/10 mL |
| Phospho-CdK1 (Tyr-15) | Cell Signalling Technology | 9111L | Rabbit | 1:1000 |
| Xenopus Cyclin B2 | Abrieu et al. 1997(24) | - | Rabbit | 2 ug/10 mL |
| phosphorylated Arpp19 (S67) | Cell Signalling Technology | 5240 | Rabbit | 1/1000 |
| Xenopus FAM122A | Lacroix et al. 2024(14) | - | Rabbit | 5 ug/10 mL |
| DyLight™ 800 anti-Rabbit | Invitrogen | A5-10036 | Goat | 1:5000 |
| DyLight™ 680 anti-Mouse | Invitrogen | 35518 | Goat | 1:5000 |
| IRDye 800 anti-Rabbit | Licor | 926-32211 | Goat | 1:10.000 |
| IRDye 800 anti-Mouse | Licor | 926-32210 | Goat | 1:10.000 |
| IRDye 800 anti-Rat | Licor | 926-32219 | Goat | 1:10.000 |
| IRDye 680 anti-Mouse | Licor | 926-68070 | Goat | 1:10.000 |

### Generation of FAM122A S158 phospho-antibody

The FAM122A S158 phospho antibody was generated by conjugating the KQCFS(p)PSLQSF peptide to KLH and injecting rabbits with the conjugate. The antibody was affinity purified against first KQCFSPSLQSF and subsequently KQCFS(p)PSLQSF. Peak fractions were kept at -80.

### Whole-cell extracts

For whole-cell extracts, cells were lysed in appropriate low salt buffer, and cell lysates were cleared by centrifugation. Protein concentration was determined using the Pierce BCA protein assay kit (Thermo Fisher Scientific) following the company’s protocol.

### Immunoprecipitation (IP)

Immunoprecipitations were performed in two conditions, transient transfection and stable cell lines. For the IP from transient transfection, HeLa cells were transfected with the indicated constructs as stated in the transfection section. After 24 hours of incubation, cells were collected, lysed in low salt buffer, and sonicated for 10 minutes (30 seconds ON/OFF cycles) at 4 °C. The lysates were then cleared by centrifugation. After centrifugation, the pellet was discarded and the supernatant was moved to new Eppendorf tubes for protein quantification. Protein concentration was determined using the Pierce BCA protein assay kit for each sample, and then, concentrations were equalized to the lowest one from the experiment. From each equalized sample. Then, proteins were purified by adding GFP-trap beads (ChromoTek) for 1 hour at 4 °C, following the company’s protocol. After IP, beads were centrifugated, and the supernatant was discarded. The beads were then washed three times in ice-cold low salt buffer before resuspending in 40 µL of NuPage LDS sample buffer.

### Western blot (WB)

Western blots were performed following standard procedures in the lab. First, samples were diluted in an appropriate volume of NuPage LDS sample buffer (Life Technologies) and were boiled for 5 minutes at 95 °C. Later, samples were analyzed by sodium dodecyl sulfate-polyacrylamide gel electrophoresis (SDS-PAGE) using a NuPage Bis-Tris 4–12% protein gel (Invitrogen). Gels were run in NuPAGE MOPS SDS Runnin Buffer (Invitrogen), and proteins were transferred to PVDF membranes (Immobilon-FL, Merck) in transfer buffer for 150 minutes at 300 mA at 4°C. After transference, membranes were blocked in 5% skim milk diluted in PBS-Tween 0.1% (PBS-T) for 30 minutes and incubated in primary antibody at 4 °C overnight. The antibodies were prepared in fresh blocking solution and used at the indicated dilutions in the antibody methods section. After primary incubation, membranes were washed three times in PBS-T 0.1% for 5 minutes each wash. Then, membranes were incubated in secondary antibody for 1 hour at room temperature and washed again. Secondary antibodies were also prepared in fresh blocking solution and used at the indicated dilutions in the antibody methods section. After the last wash, membranes were imaged and quantified using the Odyssey® CLx (LI-COR).

### Phosphatase activity assay

HeLa cells were transfected with the indicated constructs for 24 hours. After 24 hours of incubation, cells were collected (with TBS washes), lysed in low salt buffer, and sonicated for 10 minutes (30 seconds ON/OFF cycles) at 4 °C. The lysate was then cleared by centrifugation. After centrifugation, the pellet was discarded and the supernatant was moved to new Eppendorf tubes for protein quantification. Protein concentration was determined using the Pierce BCA protein assay kit for each sample, and then, concentrations were equalized to the lowest one from the experiment. Each equalized sample is split into two different Eppendorf. Then, proteins were incubated for 30 minutes at 4 °C with Protein G beads (without antibody conjugation). After incubation, samples were centrifugated and the supernatant was moved to new Eppendorf tubes, discarding the beads. B55 primary antibody was added 1:200 to one out of the two tubes from each sample, and all of the samples (with and without antibody) were incubated for 1 hour at 4 °C. After incubation, Protein G beads were added again to each sample and were incubated for 30 minutes at 4 °C. After IP, beads were centrifugated, and the supernatant was discarded. The beads were then washed twice in ice-cold low salt buffer, and one last time in activation assay buffer before eluting in 100 µL of activation assay buffer. 20 µL from each sample were taken and incubated with basic peptide (WRRApTVA)(Peptide 2.0) for 10 minutes shaking at 400 rpm at 30 °C. After incubation, PiColorLock Phosphate Detection Reagent (Abcam) was added, and samples were moved to a half-well 96-well dish, and waited 10 minutes for color development. After waiting for color development, the dish was read in FLUOstar Omega (BMG Labtech). This process was done in triplicates for each sample.

### Fluorescence-Activated Cell Sorting (FACS)

U2OS parental and U2OS FAM122A KO cell lines were seeded into 10 cm dishes. Cells were collected and resuspended in 500 µL of PBS. 1.5 mL of ice-cold 96% ethanol was added dropwise while mixing gently on a vortex and kept overnight at 4 °C. Then cells were pelleted and ethanol was discarded. Cells were washed in PBS with 1% FBS and spun down at 300g for 3 minutes. The supernatant was removed and cells were resuspended in 200 µL of Triton-X-100 0.25% in PBS and incubated for 10 minutes at room temperature. After incubation, cells were washed once in PBS with 1% FBS, resuspended in 200 µL of propidium iodide (PI) solution, and incubated overnight at 4 °C. The next day, samples were filtered through a cell strainer and analyzed in a FACS machine. The data was later analyzed with the FlowJo software and the % of cells in each cell cycle phase was determined.

### Protein purification

6xHis-Human wildtype and S158A mutant of FAM122A were produced in *Escherichia coli* and purified using TALON Superflow Metal Affinity Resin as described in (*14*)

### Interphase egg extracts

Frogs were obtained from « TEFOR Paris-Saclay, CNRS UMS2010 / INRAE UMS1451, Université Paris-Saclay», France and kept in a *Xenopus* research facility at the CRBM (Facility Centre approved by the French Government. Approval n° C3417239). Females were injected of 500 U Chorulon (Human Chorionic Gonadotrophin) and 18 h later laid oocytes were used for experiments. Adult females were exclusively used to obtain eggs. All procedures were approved by the Direction Generale de la Recherche et Innovation, Ministère de L’Enseignement Supérieur de la l’Innovation of France (Approval n° APAFIS#40182-202301031124273v4). Xenopus interphase egg extracts were obtained from oocytes arrested in metaphase II. These oocytes were treated with a Ca2+ ionophore (final concentration 2 mg/ml). Thirty minutes later, the oocytes were crushed by centrifugation twice for 20 minutes at 20,000g. The cytoplasmic fraction was recovered and used for the experiments.

### Mutagenesis

Single-point mutations of S158 of Human FAM122A was performed using Pfu ultra II fusion DNA polymerase. Oligonucleotides were purchased from Eurogentec.

See detailed Primers below.

FAM122A S158A

5’ ATT-GGG-AAG-CAG-TGT-TTT-GCG-CCA-TCC-TTG-CAA-AGT-TTT-G 3’

5’ CAA-AAC-TTT-GCA-AGG-ATG-GCG-CAA-AAC-ACT-GCT-TCC-CAA-T 3’

FAM122A S158E

5’ GGG-AAG-CAG-TGT-TTT-GAG-CCA-TCC-TTG-CAA-AG 3’

5’ CCC-TTC-GTC-ACA-AAA-CTC-GGT-AGG-AAC-GTT-TC 3’

FAM122A S158T

5’ GGG-AAG-CAG-TGT-TTT-ACG-CCA-TCC-TTG-CAA-AG

5’ CCC-TTC-GTC-ACA-AAA-TGC-GGT-AGG-AAC-GTT-TC

All other constructs were cloned by ordering the synthetic gene (Geneart) and cloning it into pcDNA5/FRT/TO YFP.

### IP-MS

Proteins were enriched from eluates using the SP3 method (*25*) and digested overnight in 25 mM ammonium bicarbonate with trypsin for mass spectrometric analysis. Digests were analyzed using an Q-Exactive Plus quadrupole Orbitrap mass spectrometer (ThermoScientific) equipped with an Easy-nLC 1000 (ThermoScientific) and nanospray source (ThermoScientific). COMET (release version 2014.01) in high-resolution mode was used to search raw data (*26*) against a target-decoy (reversed) (*27*) version of the human proteome sequence database (UniProt; downloaded 8/2020) with a precursor mass tolerance of ± 1 Da and a fragment ion mass tolerance of 0.02 D requiring fully tryptic peptides (K, R; not preceding P) and up to three mis-cleavages. Static modifications included carbamidomethylcysteine and variable modifications included oxidized methionine. Searches were filtered to a < 1% FDR at the peptide level. Quantification of LC-MS/MS spectra was performed using MassChroQ (*28*) and the iBAQ method (*29*). Missing protein abundances were imputed and bait abundances were normalized across all samples (*30*). Statistical analysis was carried out by a two-tailed Student’s t-test in Perseus

### Phosphorylation analysis of FAM122A

Proteins were enriched from eluates using the SP3 method (25), reduced and alkylated, and digested overnight in 133 mM HEPES pH 8.5. Peptides were labeled with Tandem-Mass-Tag (TMT) reagent (ThermoFisher Scientific). Each reaction was quenched by the addition of hydroxylamine to a final concentration of 0.25% for 10 minutes, mixed, acidified with TFA to a pH of about 2, and desalted over an Oasis HLB plate (Waters). The desalted multiplex was dried by vacuum centrifugation and analyzed on an Orbitrap Lumos mass spectrometer (ThermoScientific) equipped with Vanquish Neo liquid chromatography system (ThermoScientific). The raw data files were searched using COMET with a static mass of 229.162932 Da on peptide N-termini and lysines and 57.02146 Da on cysteines and a variable mass of 15.99491 Da on methionines and 79.96633 Da on serines, threonines and tyrosine against the target-decoy version of the human proteome sequence database (UniProt; downloaded 2/2013, 40,482 entries of forward and reverse protein sequences), maximum of three missed cleavages allowed, precursor ion mass tolerance 1 Da, fragment ion mass tolerance ±8 ppm, and filtered to a <1% FDR at the peptide level. Quantification of LC-MS/MS spectra was performed using in-house developed software. FAM122A phosphopeptide intensities were adjusted based on FAM122A protein abundance. Statistical analysis was carried out by a two-tailed Student’s t-test.

### AlphaFold 3 models of FAM122A-PP2A-B55

FAM122A-PP2A/B55α holoenzyme complex models were generated in AlphaFold3 (*18*) using protein sequences downloaded from Uniprot, and catalytic site metal ions available through the AlphaFold server. Seed number was set to auto. All models returned an interface predicted Template Modeling score (ipTM) and predicted Template Modeling score (pTM) above 0.5, which was used as cut-off for confidence:

FAM122A-PP2A/B55α: iPTM = 0.67 ; PTM = 0.74

FAM122A-PP2A/B55α pS158: iPTM = 0.68 ; PTM = 0.76

For each AF3 prediction, the 5 output models were aligned to assess confidence. For FAM122A-PP2A/B55 pS158 2 out of 5 predictions confidently predicted S158 in the catalytic site of PP2A C. The other 3 models had low confidence around this sequence and were not considered in our analyses.

---

**Supplementary figure 1:**
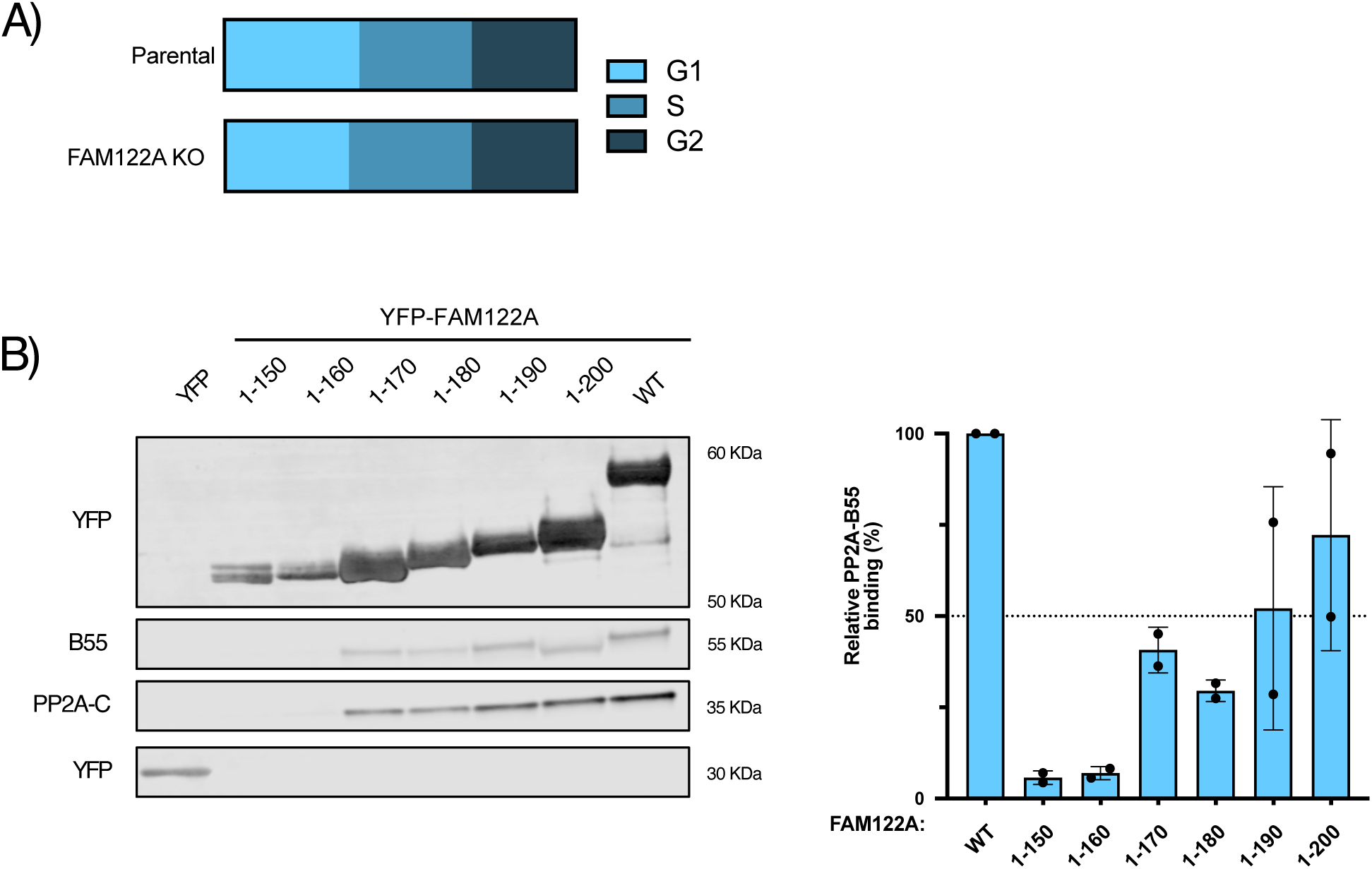
**A)** Cells from U2OS or U2OS FAM122A A2 KO cell lines were collected and analysed by Fluorescence-Activated Cell Sorting (FACS) following propidium iodide (PI) staining. Representative of three biological replicates. **B)** The indicated YFP-FAM122A tagged fragments were expressed in HeLa cells, followed by GFP trap pull down and subsequent immunoblotting of indicated proteins: PP2A-C; PP2A catalytic subunit; PP2A-B55, PP2A regulatory subunit, and YFP. Quantification showing the average and SD of two replicates assessing relative binding of FAM122A constructs indicated to PP2A-B55 normalized to WT protein.

**Supplementary figure 2:**
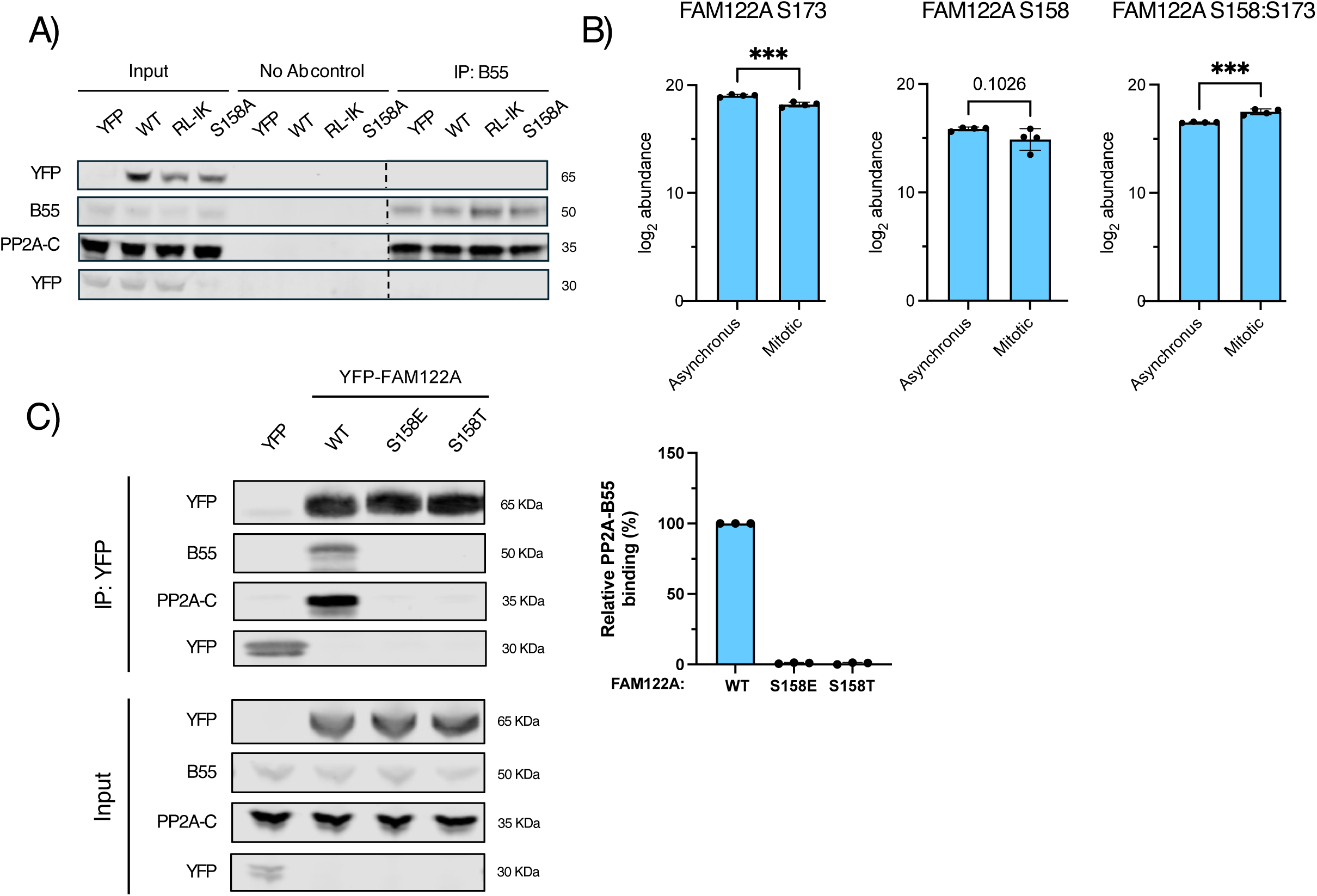
**A)** WB analysis of samples used for PP2A-B55 phosphatase assays in Fig. 4B. **B**) MS analysis of FAM122A phosphorylation in asynchronous and mitotic cells from three technical replicates. **C)** The indicated YFP-FAM122A S158 single point mutants were expressed in HeLa cells, followed by GFP trap pull down and subsequent immunoblotting of indicated proteins: PP2A-C; PP2A catalytic subunit; PP2A-B55, PP2A regulatory subunit, and YFP. Quantification of three replicates assessing relative binding of FAM122A constructs indicated to PP2A-B55 normalized to WT protein.

